# Seq2Pocket: Augmenting protein language models for spatially consistent binding site prediction

**DOI:** 10.64898/2026.01.28.702257

**Authors:** Vít Škrhák, Lukáš Polák, Marian Novotný, David Hoksza

## Abstract

Protein-ligand binding site prediction (LBS) is important for many areas of structural biology and molecular modeling, where, as in other tasks, protein language models (pLMs) have shown a great promise. In their application to LBS, the pLM classifies each amino acid as binding or not, but translating these predictions into three-dimensional binding pockets remains challenging; in particular, residue-centric predictions tend to produce spatially fragmented pockets. We present Seq2Pocket, a methodology for pocket-level LBS prediction that combines pLM finetuning, data enhancement, and structure-aware post-processing. First, we introduce sc-PDB_enhanced_, an extended training dataset that augments sc-PDB with additional small-molecule and ion-binding sites, improving coverage of non-obvious and small pockets. Second, we employ an embedding-supported smoothing classifier to refine residue-level predictions. Third, we define the Pocket Fragmentation Index and use it to select a clustering approach that preserves a consistent mapping between predictions and ground-truth pockets. We evaluate Seq2Pocket on two tasks: general binding site prediction using the LIGYSIS benchmark and cryptic binding site prediction using CryptoBench benchmark. Across both benchmarks, the proposed methodology achieves state-of-the-art performance. In particular, for general binding site prediction on the LIGYSIS benchmark, it improves distance-center-to-center recall by up to 12%, outperforming existing predictors. We believe that these findings contribute to more reliable evaluation practices in ligand binding site prediction, highlight the importance of training data curation, and provide pocket-level prediction tool that are better suited for downstream applications such as drug discovery.

## Introduction

Proteins drive most biological processes, so understanding how they work, and how to control their activity, lies at the center of fields like drug discovery, biotechnology, and agriculture. Direct experiments, whether *in vitro* or *in vivo*, take time and resources, which limits how far they can be pushed. That constraint has motivated many researchers toward *in silico* approaches, which offer a faster and more affordable way to study proteins [26].

One of the main ways to control the function of a protein is through the binding of small molecules, often called ligands. This strategy is highly successful in drug discovery, as nearly 90% of current drugs are small-molecule ligands [47]. The challenge is finding where these ligands can bind those sites. Over the years, many computational approaches have been developed to find and examine these ligand binding sites (LBS), including Molecular Dynamics (MD) simulations [11, 1], energy-based methods [35, 21, 31], docking [18], and machine learning [56, 50, 20].

Recent advances in deep learning have reshaped structural biology in two main directions. One is the emergence of structure prediction systems such as AlphaFold [27]. The other is the development of protein language models (pLM) [22, 38, 14, 23]. When combined with techniques such as transfer learning and finetuning, pLMs reach or exceed the performance of existing methods on several downstream tasks, including variant effect prediction [43], protein function prediction [52], and enzyme activity prediction [54].

Naturally, ligand binding site (LBS) prediction has also been influenced by the recent advances in deep learning [57, 37]. Since the introduction of protein language models (pLMs), a growing number of deep learning–based methods have incorporated pLM representations for LBS prediction [6, 44, 61, 19, 58]. This shift toward pLM-based approaches aligns closely with the structure of the problem: pLMs have been shown to encode local motifs [55], and binding pockets can be, to a large extend, interpreted as local motifs on the protein sequence. In typical pLM-based binding site prediction pipelines (i.e., *transfer learning* through embeddings, or *finetuning*), the model produces a per-residue score that reflects the probability of that residue belonging to a binding site. Despite the competitive performance reported for many of these approaches, a recent independent evaluation suggests that traditional methods continue to perform strongly in LBS prediction, in some cases matching or outperforming newer deep learning–based techniques [50].

When comparing the residue-level and pocket-level performance, we can find a conflict between these two metrics: while residue-level metrics might look promising, especially in cases where a large binding site provides many true positive residues, the pocket-level metrics could be disappointing. For example, if a protein has three distinct binding pockets and a predictor only finds the largest one, its pocket-level recall is only 33%, even if its residue-level scores are high.

Therefore, while residue-level performance is a prerequisite of a good predictor, it is insufficient for binding site discovery evaluation. On that account, pocket-level metrics such as distance-center-to-center (**DCC**) have become the evaluation standard [50, 32].

In this study, we identify three challenges related to LBS evaluation and the use of pLMs:

1. **Spatial incoherence**: A truly robust LBS predictor should identify continuous binding regions rather than scattered residues. We observe a tendency in pLMs to produce what we define as **incomplete pockets**: residue-wise predictions that often achieve high statistical scores but fail to form continuous binding regions (see examples in Figure 1, Figure 2 and Figure S2a, Figure S3a in the Supplementary Material).
2. **Bias towards large binding sites:** Current literature suggests that most predictors excel at identifying large orthosteric sites while struggling with smaller more challenging targets [50]. We argue that this performance issue is largely driven by limitations in existing training sets. We further show that augmenting the training data with more diverse binding conformations significantly improves the detection of these non-obvious sites.
3. **Metrics inflation via clustering fragmentation**: Mapping residue-level predictions to three-dimensional pockets requires a clustering step. We demonstrate that the choice of clustering significantly impacts pocket-level results; while one approach may degrade performance, another can paradoxically make a model appear superior by inflating pocket-level metrics like DCC. This occurs when the algorithm partitions a single binding region into numerous small clusters: a phenomenon we refer to as **clustering fragmentation**. While such fragmentation may improve numerical scores (DCC), it does not correspond to better identification of functional binding sites and reduces the practical usefulness of the predictions.

**Fig. 1.**
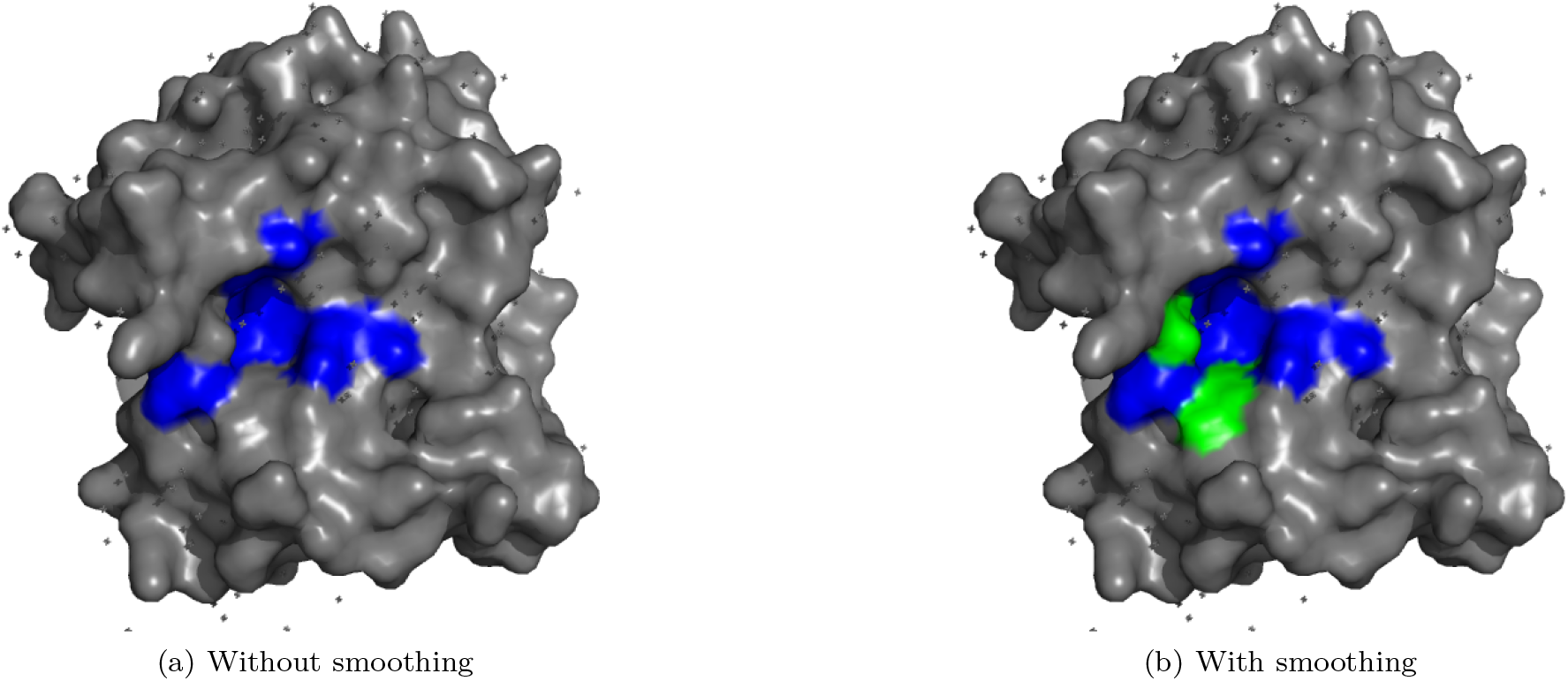
Example of residues added by smoothing classifier (GREEN: Pro8, Phe56, Ile58, Val166) of a predicted pocket (BLUE). This is a 6isuA structure.

**Fig. 2.**
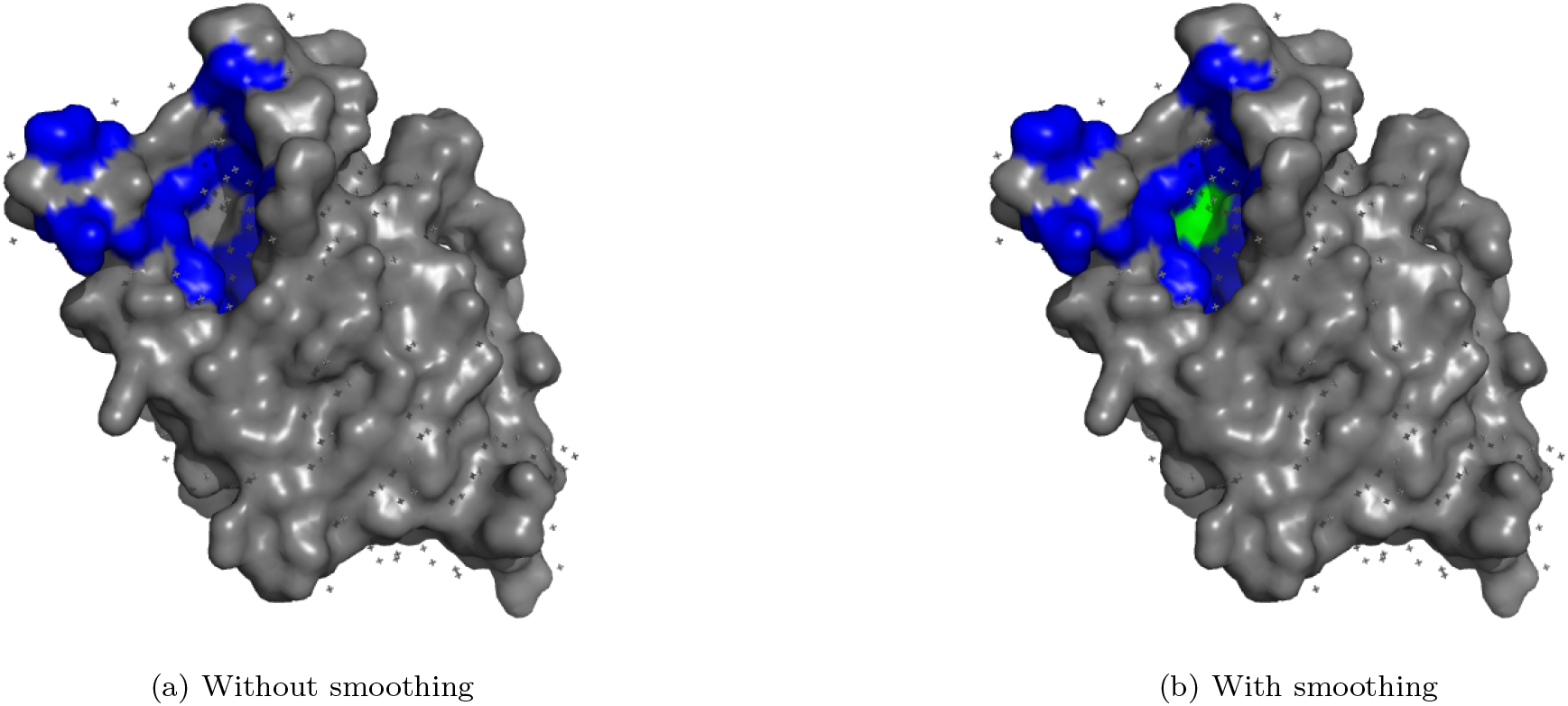
Example of residues added by smoothing classifier (GREEN: Thr1042, Thr1067) of a predicted pocket (BLUE). The cavity was not fully predicted by the finetuned pLM. Smoothing classifier fills the gap.

To address these challenges, we develop a prediction methodology named **Seq2Pocket** built upon a finetuned ESM-2 pLM, that serves as the methodology’s prediction core. This methodology contributes in three ways. First, we address the problem of **incomplete pockets** by introducing an embedding-supported smoothing classifier that uses latent representations of neighboring residues to refine residue-level predictions, ensuring structurally complete binding pockets (see Pocket Smoothing). Second, to improve generalization toward non-obvious targets, we construct **sc-PDB**_**enhanced**_ dataset that incorporates *small-molecule-binding* and *ion-binding* sites in order to reduce their omission during inference (see Datasets). Third, we define the **Pocket Fragmentation Index** (see Evaluation Framework), and conduct a comparison of different clustering strategies.

We evaluate our approach on two challenges: the prediction of general binding sites (GBS) and the prediction of cryptic binding sites (CBS). In both challenges, we were able to secure state-of-the-art performance and named the prediction methodology **Seq2Pocket**.

## Methods and Materials

Our methodology consists of a multi-stage pipeline. At its core is a finetuned ESM-2 (3B) model that serves as the residue-level prediction engine. For GBS prediction, we train two variants of this core to quantify the impact of training data quality: one on the standard sc-PDB dataset [12] and another on our augmented sc-PDB_enhanced_ (Datasets). For CBS prediction, we utilized an existing model finetuned on the CryptoBench training set [58]. Predicted residue scores are then refined using the embedding-supported smoothing classifier. Finally, spatial clustering is applied to group residues into binding pockets.

### pLM Finetuning

The GBS prediction task required finetuning of the ESM2-3B protein language model on the scPDB dataset. We first adapted the model architecture by removing the masked language modeling head and replacing it with a binary classification head operating at the residue level. Training was performed using binary cross-entropy loss with the AdamW optimizer and L2 regularization. A dropout rate of 0.3 [25] was applied, and the learning rate was set to 1e-4. Due to GPU memory constraints, the batch size was fixed to one. The model was finetuned for a total of 3 epochs, with class weighting employed to stabilize the imbalance between binding and non-binding residues (9.40% of positives in the LIGYSIS dataset). The finetuning was performed in two stages. In the first stage, the weights of the entire pLM backbone were frozen, allowing only the newly initialized binary classification head to be trained. In the second stage, the entire model (backbone and head) was unfrozen and trained jointly. The threshold for assigning the positive class was set to 0.7. All parameters were set based on a validation run using one of the CryptoBench train splits as a validation set.

The whole architecture is depicted in Figure S5 in the Supplementary. We also finetuned the 650M variant of ESM2 for availability on hardware with restricted resources.

For the CBS task, a finetuned ESM2-3B model from a recent study [58] was employed without any adjustments.

### Pocket Smoothing

We introduce a binary classifier designed to decide whether a candidate residue should be incorporated into a predicted binding pocket. The purpose is to recover residues that are missed in the initial residue-level prediction.

The classifier is implemented as a three-layer feed-forward neural network. It consists of two hidden layers with 2048 neurons each, followed by a single-neuron output layer with a sigmoid activation for binary classification. Training is carried out for 19 epochs using a batch size of 512 and a dropout rate of 0.5. We optimize the model using binary cross-entropy loss with the AdamW optimizer and L2 regularization. Class weighting is applied to counter the imbalance between positive and negative examples. Hyperparameters are selected based on validation experiments on one of the training subsets, optimizing the area under the precision–recall curve (AUPRC).

Each input to the classifier is formed by concatenating two embeddings: an embedding of the candidate residue and a pocket context embedding. Both are taken from the final hidden layer of the ESM-2 (3B) protein language model.

The pocket context embedding is intended to represent the local environment of a predicted binding site. For a given candidate residue, we identify all neighboring residues within a 15 Å radius that were initially predicted as binding residues by the pLM. The embeddings of these neighboring residues are then aggregated using mean pooling to produce the pocket context embedding.

The classifier is trained using annotated binding sites from the CryptoBench dataset. Training samples are constructed as follows:

- **Positive Examples:** residues that belong to a known binding site.
- **Negative Examples**: non-binding residues located within 10 Å of a binding site.

During inference, a negatively predicted residue is considered for smoothing if it lies within 15 Å from any positively-predicted residue.

The geometric thresholds were determined using one of the CryptoBench training splits as a validations set.

An overview of the pocket smoothing architecture is provided in Figure S6 in the Supplementary Material.

### Clustering

After residue-level prediction and smoothing, predicted binding residues are grouped into discrete three-dimensional regions using a surface-based clustering procedure. For each atom belonging to a predicted binding residue, we generate solvent-accessible surface (SAS) points using the Shrake–Rupley algorithm [45], with a probe radius of 1.6 Å and a point density of 50. We employ a modified implementation from the Biopython library [10]. The resulting SAS point cloud is then clustered into candidate binding pockets using the Mean Shift algorithm [7] with a bandwidth of 9 Å. An example of the resulting SAS point distribution for a correctly predicted binding site on human C-terminal Src kinase (PDB ID: 1BYG, chain A [34]), where the Staurosporine inhibitor binds, is shown in Figure S1 in the Supplementary Material.

Each surface-based cluster was then mapped back to residues using a voting scheme: each solvent-accessible surface (SAS) point inherits the cluster assignment produced by the Mean Shift algorithm, and each residue contributes multiple SAS points. A residue is assigned to the cluster for which the majority of its SAS points vote. Once residues are assigned to clusters, each resulting pocket is scored by computing the mean predicted binding probability of its residues, as provided by the finetuned pLM. These pocket scores are subsequently used to rank the predicted binding sites.

## Datasets

This work utilizes three datasets (Table 1):

- **sc-PDB** for training the GBS predictor,
- **LIGYSIS** for evaluating the GBS predictor,
- **CryptoBench** for evaluating the CBS predictor.

**Table 1.** Datasets statistics. LIGYSIS and CryptoBench test sets were used for evaluation, sc-PDB and CryptoBench train sets were used for training. Although the sc-PDB and sc-PDB_enhanced_ datasets contain the same structures, the sc-PDB_enhanced_ contains additional pockets found either in that structure or in a different conformation of the same protein.

| Dataset | Purpose | # of structures | # of residues | # of binding residues | % of binding residues | # of pockets |
| --- | --- | --- | --- | --- | --- | --- |
| LIGYSIS | GBS evaluation | 2,775 | 832,071 | 78,205 | 9.40% | 6,882 |
| sc-PDB | GBS training | 17,077 | 5,558,460 | 536,011 | 9.64% | 17,077 |
| sc-PDB<sub>enhanced</sub> | GBS training | 17,077 | 5,558,460 | 824,046 | 14.83% | 56,968 |
| CryptoBench test subset | CBS evaluation | 184 | 55,572 | 3,158 | 5.68% | 215 |
| CryptoBench train subset | CBS training | 762 | 233,435 | 13,342 | 5.72% | 872 |

### LIGYSIS Dataset

The LIGYSIS dataset is the most recent dataset of ligand binding sites. It contains non-redundant protein-ligand interfaces. To remove redundancy, as well as take advantage of all available data, the annotations were mapped to a single representative biological unit using UniProt mapping. The LIGYSIS also includes ion binding sites; in fact, about 20% of the pockets are binding ions. We used its subset of 2775 human protein chains, which was utilized in a recent study to evaluate the performance of 11 ligand-binding site predictors [49]. This allowed us a direct comparison with these evaluated methods.

### sc-PDB Dataset

The sc-PDB dataset is one of the most widely used datasets for training binding site prediction methods [44, 2, 46, 29, 28]. We used sc-PDB v2017, the most recent version, which contains 17,077 structures^1^. sc-PDB has been reported to have 9.7% overlap with the LIGYSIS dataset [50]. Unfortunately, the original sc-PDB dataset considers only the most relevant ligand for each entry. This is not biologically accurate because a single protein can contain more than one binding site. Furthermore, sc-PDB does not consider ion binding sites. Therefore, we subsequently extended the set of sc-PDB binding sites by using AHoJ-DB [16, 17] (version *v2c*). We selected AHoJ-DB structures that were also present in the sc-PDB and added AHoJ-DB binding sites that were not found in the sc-PDB, including ion binding sites. This expansion increased the total number of pockets from 17,077 to 56,968 (Table 1). We refer to this version as sc-PDB_enhanced_.

We note that the reported 9.7% pocket overlap between sc-PDB and the LIGYSIS test set is problematic. This overlap also affects sc-PDB_enhanced_, since its additional pockets are derived from the same underlying protein structures. Therefore, in any subsequent training, we removed the overlapping UniProt sequences from the training data in both sc-PDB and sc-PDB_enhanced_. As a result, the number of protein structures used for training was reduced from 17,077 to 13,140. Correspondingly, the total number of annotated pockets was reduced from 17,077 to 13,140 in sc-PDB and from 56,968 to 38,578 in sc-PDB_enhanced_. All training experiments reported in this work use these filtered datasets. Pocket counts reported in Table 1 refer to the unfiltered datasets and are provided for reference.

### CryptoBench Dataset

The CryptoBench dataset [60] is built on top of AHoJ-DB [16, 17] via a multi-stage processing pipeline, including structural quality check filtering (TM-score ≥ 0.5), resolution filtering (≤ 2.5 Å) and ensuring geometric compactness of pockets by controlling the radius of gyration (max. 20% change). Most importantly, it applies a crypticity criterion based on pocket RMSD. Specifically, the pocket is marked as cryptic, if there exists a conformation where the RMSD of the pocket atoms between ligand-bound state (holo) and the ligand-free state (apo) exceeds 2 Å. To remove redundancy, the resulting structures are filtered down to 40% sequence identity [48]. The dataset provides pre-defined training and test splits: to ensure no information leakage occurs between them, we applied a secondary filter to the remaining sequences once again, this time to 10% sequence identity, meaning that no two sequences from different subsets share more than 10% sequence identity. It should be noted that CryptoBench does not include ion binding sites. In this work, we used a subset of CryptoBench that excludes binding sites spanning multiple chains.

### Evaluation Framework

Our evaluation uses two benchmarks: the LIGYSIS dataset [50] for GBS and CryptoBench [60] test set for CBS. Each example in these datasets can be derived from multiple structural sources. For LIGYSIS, binding site annotations are often aggregated from several PDB structures; we follow the dataset’s convention by utilizing the predefined representative structure and chain ID during the evaluation. In the case of CryptoBench, binding sites are provided in both apo (ligand-free) and holo (ligand-bound) forms [5]. We perform our evaluation exclusively on the apo structures, as this represents the most relevant scenario for the discovery of the binding site, where the bound conformation is unknown and the predictor must identify the site on an unbound protein surface.

We report standard classification metrics at the residue-level: Matthews Correlation Coefficient (MCC), Area Under the ROC Curve (AUC), Area Under the Precision-Recall Curve (AUPRC), F1-score, and Accuracy (ACC). As the datasets are heavily imbalanced (only 5% are positive examples in some datasets), MCC, AUPRC, and AUC serve as more robust performance indicators. While ACC and F1-score are reported for completeness, they are less optimal for highly imbalanced datasets [8].

The main focus of this study is on pocket-level metrics, specifically **DCC-based (distance-center-to-center) recall** and **Relative Residual Overlap** (**RRO**). DCC measures the distance between the geometric centers of the predicted binding site and the ground-truth binding site. A prediction is a ‘hit’ if the distance between the predicted and ground-truth pocket centers is **below 12 Å** [50].

To test how well the model ranks its predictions, we calculate recall using different selection limits. For a protein with *N* experimentally known sites **DCC**_**top-N**_ considers only the *N* highest-scoring predicted pockets. However, protein datasets are often sparsely annotated; a model might correctly identify a valid but unannotated site as its top prediction, which would push a known site out of the list of top-*N* results. To account for this, we report **DCC**_**top-(N+K)**_, where we set K=2 to provide a budget of two additional predictions. This metric assesses whether the N ground-truth pockets are successfully identified within the model’s top N+K candidates. Finally, **DCC**_**MAX**_ considers all predicted pockets regardless of N; while this provides an upper bound for recall, it should be interpreted alongside precision to avoid favoring models that over-predict or over-fragment their predictions.

Next, RRO was computed for each pocket from the evaluation dataset as

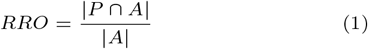

where *P* represents the number of residues from a predicted pocket and *A* represents the number of residues from the actual pocket.

To evaluate the clustering method, we suggest a measure called Pocket Fragmentation Index (PFI), which measures the average number of predicted clusters assigned to each ground-truth pocket:

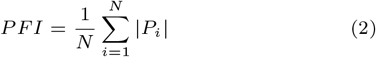

where *N* is the total number of ground-truth pockets and |*P*_i_| is the number of predicted clusters that overlap the i-th pocket. An ideal clustering strategy achieves a one-to-one mapping, resulting in a PFI of 1.0. By monitoring this index, we can identify **excessive clustering fragmentation**, where a single biological binding site is incorrectly partitioned into multiple predicted regions. A high PFI indicates that, while the residues may be correctly predicted, the structural representation is partitioned into too many segments, making later downstream applications more difficult. Partitioning the binding site into more fragments can also artificially help inflating the DCC-based metrics.

Although cryptic binding sites show structural flexibility, they typically exhibit localized conformational changes, such as loop rearrangements or secondary structure displacements, rather than a total pocket reorganization [4, 51]. Therefore, the same structural metrics (DCC-based recall, coverage, and PFI) remain applicable for the CBS task.

The evaluation framework for the GBS task is identical to that of Utgés et al. [50], which allows direct comparison between our results and other state-of-the-art methods measured on the LIGYSIS dataset. We report only the best-performing methods for comparison in both residue-level and pocket-level metrics. Similarly, for the CBS task, the performance at the residue-level of the transfer learning baseline is sourced from Škrhák et al. [58], as the study uses the same evaluation dataset.

## Results

We conducted an ablation study for both the GBS and CBS models by evaluating each in two configurations: without pocket smoothing and with the proposed pocket smoothing classifier.

### Residue-level Evaluation

Table 2 summarizes the residue-level performance in the LIGYSIS and CryptoBench evaluation sets. Although Accuracy (ACC) and F1-score are reported for completeness, the main focus of this evaluation is on MCC, AUC, and AUPRC, as these metrics are more robust to class imbalance.

**Table 2.** Residue-level performance comparison across GBS and CBS tasks. The table compares our proposed finetuning and smoothing approach against state-of-the-art baselines using residue-level metrics. Performance metrics for *Baseline transfer learning, PUResNet, and P2Rank*_CONS_are adopted from existing benchmarks [50, 58]. Note that for the GBS task, only the top-performing methods from the LIGYSIS study [50] are included for comparison. Primary emphasis is placed on MCC, AUC, and AUPRC; note that AUC and AUPRC are invariant to the smoothing process as it modifies binary labels without altering the underlying probability distributions. As *PUResNet* and *fpocket* don’t predict residue-level probabilities, AUC and AUPRC could not be reported. F1-scores and Accuracy (ACC) for PUResNet and P2Rank_CONS_were independently calculated using our evaluation framework, specifically utilizing weighted averaging for F1-scores to better account for class imbalance.

| Evaluation | Method | Train set | AUPRC | AUC | MCC | ACC | F1 |
| --- | --- | --- | --- | --- | --- | --- | --- |
| CryptoBench<br>test set<br>(CBS) | Seq2Pocket | CryptoBench train | **0.475** | **0.894** | 0.460 | 92.4% | 0.932 |
|  | Seq2Pocket (without smoothing) | CryptoBench train | **0.475** | **0.894** | **0.473** | 93.5% | 0.939 |
|  | *CryptoBench baseline* [58] | CryptoBench train | 0.382 | 0.871 | 0.395 | 93.1% | 0.933 |
| LIGYSIS<br>(GBS) | Seq2Pocket | sc-PDB<sub>enhanced</sub> | **0.552** | **0.853** | 0.438 | 87.2% | 0.884 |
|  | Seq2Pocket (without smoothing) | sc-PDB<sub>enhanced</sub> | **0.552** | **0.853** | **0.468** | 89.4% | 0.899 |
|  | Seq2Pocket | sc-PDB | 0.490 | 0.816 | 0.410 | 88.4% | 0.889 |
|  | *PUResNet* [50, 29] | sc-PDB | - | - | 0.39 | 92.7% | 0.917 |
|  | *P2Rank<sub>CONS</sub>* [50, 32, 33] | CHEN11 | 0.46 | 0.76 | 0.30 | 85.9% | 0.870 |
|  | *fpocket* [50] | - | - | - | 0.12 | 52.4% | 0.616 |
|  | *Ligsite* [50, 24] | - | 0.38 | 0.70 | 0.21 | 74.7% | 0.795 |

The results show that pLM finetuning represents a state-of-the-art strategy, as our finetuned models surpass all other reported methods across both datasets. Specifically, the GBS models outperform the second-best reported method by more than 9% in both AUC and AUPRC, and by 13% in MCC (Table 2). Nevertheless, we observe that the application of the pocket smoothing classifier leads to a modest decrease in MCC.

### Smoothing and Pocket-level Evaluation

As summarized in Table 3, pLM finetuning shows improved performance in DCC_top-N_, DCC_TOP-(N+2)_ and RRO over the fpocket rescored by PRANK (denoted fpocket_PRANK_), setting a new state-of-the-art performance on the LIGYSIS dataset. In particular, our finetuned approach shows a substantial improvement of 12% in DCC_top-N_ recall and 8% in DCC_top-(N+2)_ recall compared to previous state-of-the-art (fpocket_PRANK_, P2Rank_CONS_). Applying pocket smoothing further increases DCC recall, adding on average 8.62 residues per structure (median: 6.0)^2^. While Ligsite achieves the highest RRO among the tested methods, its DCC recall remains limited.

**Table 3.** Pocket-level performance comparison across GBS and CBS tasks. The table compares our proposed finetuning and smoothing approach against state-of-the-art baselines using pocket-level metrics. The top-performing methods in pocket-level metrics from the LIGYSIS study are added, namely fpocket [36] and Ligsite [24]. The emphasis is on DCC_top-N_ and DCC_top-(N+2)_, which don’t reward excessive overprediction (i.e., predicting too many binding sites).

| Evaluation | Method | Train set | DCC<sub>top-N</sub> | DCC<sub>top-(N+2)</sub> | DCC<sub>MAX</sub> | % RRO |
| --- | --- | --- | --- | --- | --- | --- |
| CryptoBench<br>test set<br>(CBS) | Seq2Pocket | CryptoBench train | **66.1%** | **76.3%** | **76.7%** | **52.9%** |
|  | Seq2Pocket (without smoothing) | CryptoBench train | 64.7% | 74.9% | 74.9% | 49.2% |
|  | *CryptoBench baseline* [58] | CryptoBench train | 61.9% | 73.0% | 75.3% | 42.1% |
| LIGYSIS<br>(GBS) | Seq2Pocket | sc-PDB<sub>enhanced</sub> | **61.0%** | **68.8%** | 72.3% | 52.9% |
|  | Seq2Pocket (without smoothing) | sc-PDB<sub>enhanced</sub> | 60.4% | 66.8% | 68.7% | 52.9% |
|  | Seq2Pocket | sc-PDB | 51.0% | 57.3% | 59.4% | 49.6% |
|  | *PUResNet* [50, 29] | sc-PDB | 40.6% | 41.1% | 41.1% | 61.0% |
|  | *P2Rank<sub>CONS</sub>* [50, 32, 33] | CHEN11 | 48.8% | 53.9% | 57.0% | 56.4% |
|  | *fpocket<sub>PRANK</sub>* [50] | - | 48.8% | 60.4% | **91.3%** | 52.6% |
|  | *Ligsite* [50, 24] | - | 41.3% | 48.4% | 49.7% | **77.6%** |

The model trained on the original sc-PDB showed 48.3% DCC_top-(N+2)_ on small binding sites (less than 10 residues) and 74.1% on large binding sites (10 or more residues). In comparison, after enhancing the training set, the model trained on sc-PDB_enhanced_ achieves 59.3% on small and 82.0% on large binding sites. Therefore, dataset enhancement led to 11% improvement on small binding sites and 8% on large binding sites (see Table S1 in Supplementary Material for more details).

Lastly, We utilized P2Rank [32], a state-of-the-art geometry-based predictor. P2Rank is a widely used tool in the LBS task. It identifies binding pockets by analyzing the local geometry of the protein surface instead of relying on sequence embeddings. Interestingly, although P2Rank was not trained to reveal cryptic binding sites, it showed competitive performance on the CBS task in multiple studies [13, 60]. We run P2Rank on the CryptoBench test set and merged the predictions with the output of the smoothed pLM. The merged prediction achieved DCC_MAX_ greater than 80% and a mean RRO of 59% (see Figure S7 in Supplementary).

Comparing the finetuned pLM with the CryptoBench baseline reveals comparable DCC scores but a significant increase in RRO (Table 3). This indicates that the pLM identifies more complete binding pockets than the CryptoBench baseline.

To make the models available for hardware settings with more constrained computational resources, we repeated finetuning for the ESM-2 (650M) variant, and compared the performance with the ESM-2 (3B) models (see Table S1 in Supplementary Material.)

### Selecting an Optimal Clustering Strategy

To select the optimal clustering strategy, we evaluated candidate methods based on DCC-based recall and the **Pocket Fragmentation Index** (PFI). An ideal approach identifies binding sites with high localization accuracy (DCC) while maintaining structural integrity through low PFI.

Our analysis performed on the LIGYSIS dataset shows that relying solely on DCC can be misleading. As illustrated in Figure 3, some methods like DBSCAN [15] achieve high pocket-level performance (DCC_top-(N+2)_ = 69.2%) only because they fragment a single pocket into numerous small clusters (PFI = 4.68). Although adjusting the DBSCAN parameters can reduce this fragmentation, it simultaneously leads to a significant drop in DCC recall (DCC_top-(N+2)_ = 46.4%, PFI = 1.46). Our clustering approach, proposed in Methods and Materials, exhibits low fragmentation (PFI = 2.12) while maintaining high pocket-level performance (DCC_top-(N+2)_ = 68.8%). All tested configurations are detailed in the Supplementary Materials, section 1.

**Fig. 3.**
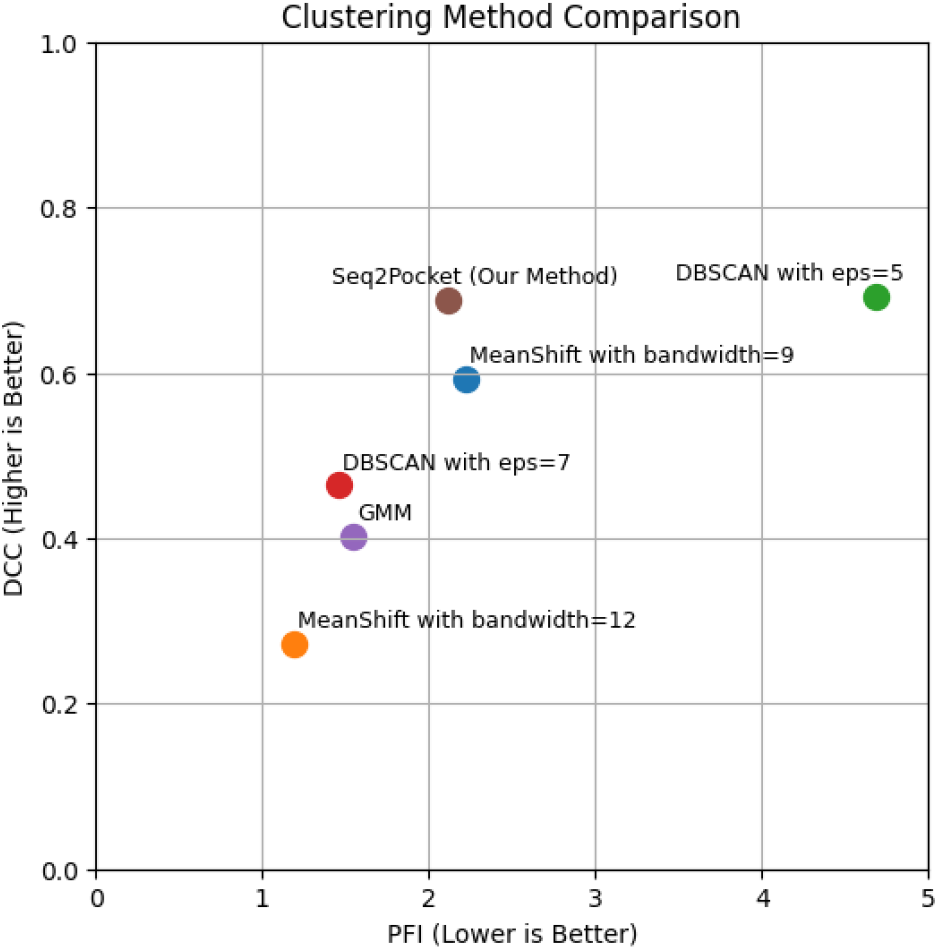
How to select an optimal clustering method. Figure reports the DCC_top-(N+2)_ metric on the LIGYSIS benchmark. The optimal method should be located in the top-left sector, i.e., showing strong DCC performance while maintaining low fragmentation. All tested configurations and their parameters are detailed in the Supplementary Materials, section 1.

## Discussion

One of the most interesting observations made during the development of the Seq2Pocket model is that the performance of LBS predictors is strongly influenced by factors beyond the core model architecture. In particular, we show that **training data** and **post-processing** play an important role in binding site prediction performance.

### Bias towards large binding sites

Current benchmarks and datasets are heavily biased toward large, orthosteric binding sites, which are overrepresented in datasets such as sc-PDB. As a result, many predictors are underperforming on smaller, less obvious sites. This limitation is not primarily a modeling issue, but rather a consequence of incomplete training data. Many datasets frequently include only the main ligand binding sites (often just the primary orthosteric site) and ignore additional small-molecule or ion-binding events, or binding sites observed in alternative conformations of the same protein.

We reduce this bias by constructing the **sc-PDB**_**enhanced**_ dataset and training our model on it, which incorporates all experimentally observed binding sites including ions. The resulting model achieves improvements across all evaluated metrics, outperforming the model trained on the original sc-PDB by more than 5% in AUPRC, and 10% in DCC_top-N_ at the pocket level (see Table 3). These results confirm that augmenting training data with diverse binding conformations is essential for improving generalization toward non-obvious targets.

We also note that while traditional datasets like sc-PDB focus solely on organic drug-like ligands [30, 41, 12, 60, 40, 9, 50], LIGYSIS treats also ions as valid binding molecules. There is no universal “correct” definition; rather, the inclusion of ions depends on the specific downstream application. However, this needs to be taken into account when collecting training data. As demonstrated in Table 3, the improvement from enhancing the dataset is significant (i.e., *>*10% in DCC). Therefore, we strongly encourage developers of binding site prediction methods not only to rely on the pre-existing binding site datasets but also to screen other available protein structures [3] for additional binding sites that might be included into their training sets, for example, through sequence and structure similarity.

### Metrics inflation through clustering fragmentation

Translating probabilities into meaningful 3D pockets is sensitive to the clustering strategy. We demonstrate that high pocket-level recall, particularly DCC, can be artificially inflated through **clustering fragmentation**, where a single functional binding region is partitioned into multiple small predicted pockets. While this fragmentation increases the chance that at least one cluster lies close to the pocket center, it does not improve the identification of binding sites and substantially reduces the practical usability of the predictions.

We therefore define **Pocket Fragmentation Index**, which measures the correspondence between predicted and actual pockets. Some methods such as DBSCAN often achieve high recall by producing severely fragmented outputs (PFI *>* 4). Our proposed pipeline maintains a reasonable balance between pocket-level performance and binding site integrity.

### Spatial incoherence

To recover residues initially missed by the finetuned pLM during prediction (as illustrated in Figure 1, Figure 2 and Figure S2, Figure S3 in the Supplementary Material), we perform additional post-processing step that leverages surounding residues context. Smoothing improves not only the visual completeness of predicted pockets, but also pocket-level metrics such as DCC and RRO.

### Implications for evaluation practice

Taken together, our results show that strong residue-level performance does not guarantee accurate pocket-level predictions. Several recent studies report only residue-level metrics and omit pocket-level evaluation altogether [42, 39, 53], which can delude actual pocket level performance. As demonstrated by the disagreement between Table 2 and Table 3, evaluating both levels is essential. Furthermore, we recommend controlling for artificial performance inflation at the pocket level, for example by reporting fragmentation-aware measures such as the proposed Pocket Fragmentation Index (PFI).

## Conclusion

Protein language models are becoming essential in computational biology pipelines. This work addresses the challenge of transitioning from sequence-level pLM predictions to 3D structural pockets. We demonstrate that the inherent nature of sequence-level predictions can lead to incomplete binding sites, a problem we mitigate through a smoothing classifier and a carefully selected clustering strategy. By combining these refinements with high-quality training data, our framework surpasses state-of-the-art methods on two major benchmarks: CryptoBench for cryptic pocket prediction and LIGYSIS for general pocket detection. We believe that our insights will stimulate further development in the binding site prediction domain.

## Author contributions statement

D.H. and M.N. conceived the research, V.S., L.P. and D.H. conceived the experiments, V.S. and L.P. conducted the experiments, V.S. wrote the manuscript, all authors discussed the results and reviewed the manuscript.

## Data and code availability

The Seq2Pocket web interface is available at https://seq2pocket.projekty.ms.mff.cuni.cz.

The code created for the purpose of this manuscript can be found at https://github.com/skrhakv/seq2pocket. The repository also includes a short tutorial for running the proposed methodology. The sc-PDB_enhanced_ is published using the Zenodo platform at https://zenodo.org/records/18271517 [59]. The trained models, dataset annotations, and prepared PyMOL scripts for visualization of the predictions can be downloaded at https://s3.cl4.du.cesnet.cz/a93fcece52e6da0dd335b4459d47b0aebb74836b:share/seq2pocket.tar.gz.

## Acknowledgments

This work was supported by the Czech Science Foundation (GAČR) grant number 23-07349S, the ELIXIR CZ Research Infrastructure (ID LM2018131, MEYS CR), the Charles University Grant Agency (GAUK) project number 260 125 and the Charles University project SVV number 260 821. Computational resources were provided by the e-INFRA CZ project (ID:90254), supported by the Ministry of Education, Youth and Sports of the Czech Republic.

## Footnotes

1 The whole dataset contains 17,594 structures, but due to annotation mismatches, we were able to retrieve only 17,077 structures

2 Measured on Seq2Pocket that was trained on scPDB_enhanced_

