## supplementary for "Seq2Pocket: Augmenting protein language models for spatially consistent binding site prediction"

#### This PDF file includes:

Supporting text
Figs. S1 to S7
Tables S1 to S2
SI References

### Supporting Information Text

#### 1. Tested configurations for clustering

In addition to the clustering method described in the main text, we evaluated three alternative configurations. Although our primary approach clusters Solvent Accessible Surface (SAS) points, these alternative configurations operate by clustering residues. We utilized several algorithms from the *scikit-learn* library, specifically DBSCAN ($eps \in \{5, 7\}$, where $eps$ is maximum distance between two points to be considered part of a single cluster), MeanShift ($bandwidth \in \{9, 12\}$), and Bayesian Gaussian Mixture Modeling (GMM). The evaluation workflow was structured as follows:

- Extraction of predicted binding residues using pLM.
- Clustering of identified residues using one of the candidate algorithms.
- Application of a smoothing classifier to refine the spatial boundaries of the predicted pockets.

Although the implementation of DBSCAN and MeanShift followed standard procedures (i.e., initializing the sklearn algorithm object and executing $fit\_predict$ method), the Bayesian Gaussian Mixture Modeling (**GMM**) approach required a two-stage process:

1. **Cluster Estimation:** An initial GMM run was performed to estimate the optimal number of clusters. A cluster was considered significant if it contained at least 10% of the total predicted binding residues.
2. **Final Clustering:** The Bayesian GMM was re-initialized and executed using the number of significant clusters determined in the first stage.

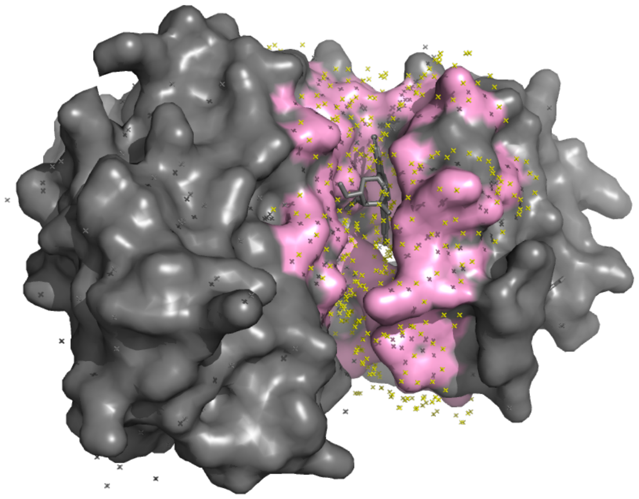

**Fig. S1. SAS points on predicted residues.** Here, the fine-tuned GBS-pLM model correctly predicts a binding site on the human C-terminal Src kinase (*PDB ID: 1BYG, chain A*), where an inhibitor Staurosporine binds. The yellow dots represent the solvent accessible surface (**SAS**) points, that are subsequently clustered using MeanShift algorithm (1).

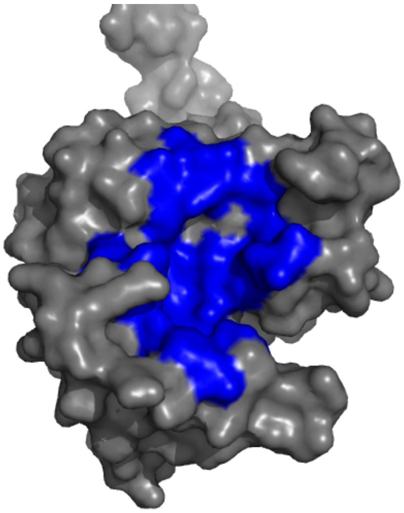

**(a)** Without smoothing

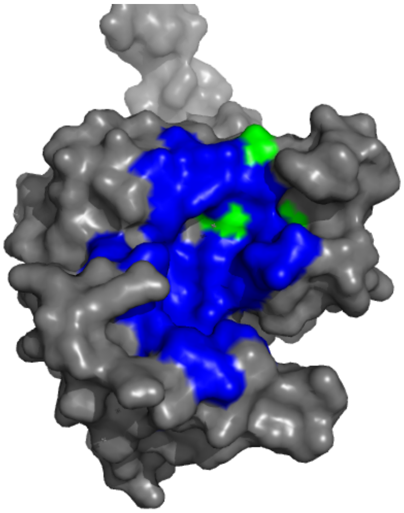

**(b)** With smoothing

**Fig. S2.** Example of residues added by smoothing classifier (GREEN: Thr213, Asp216, Leu218) of a predicted pocket (BLUE). This is a 7nlxA structure from the CryptoBench test set (2).

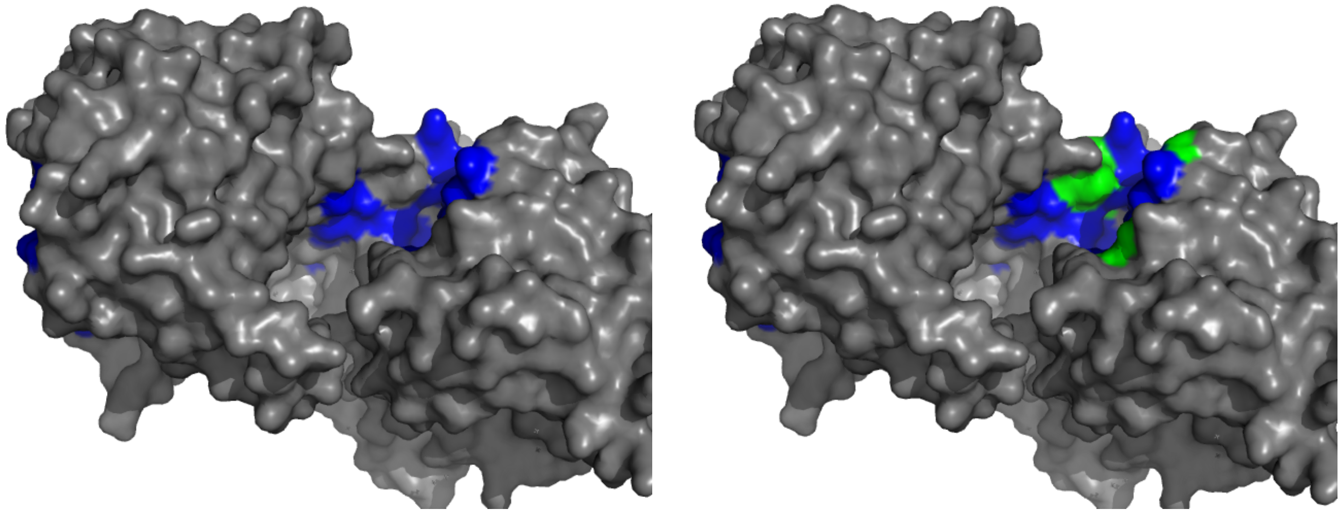

**(a)** Without smoothing **(b)** With smoothing

**Fig. S3.** Example of residues added by smoothing classifier (GREEN: Gly25, Thr63, Phe65, Ala112, Met300, Phe304, Phe305) of a predicted pocket (BLUE). This is a 7o1iA structure from the CryptoBench test set (2).

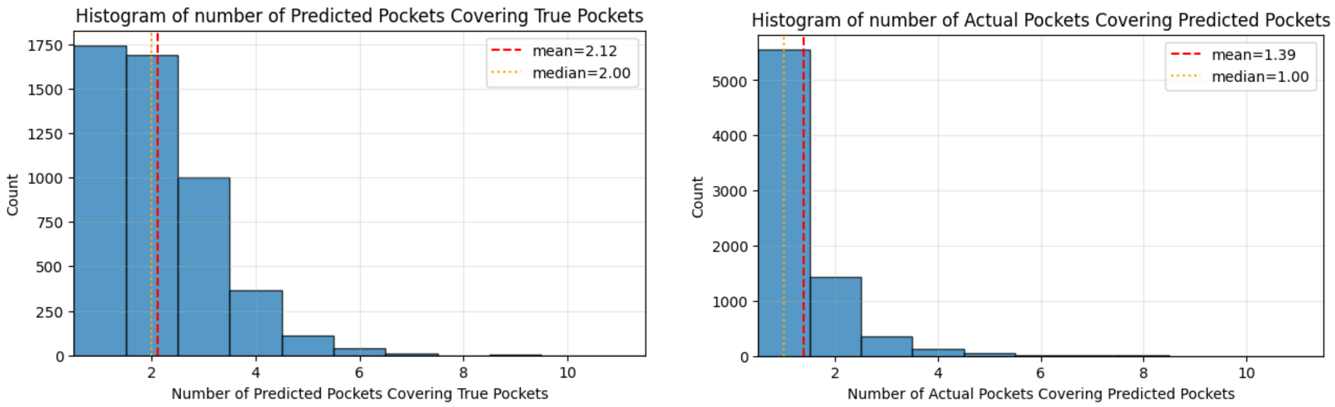


**Fig. S4. Evaluation of predicted vs. actual pocket mapping.** The **left diagram** shows the average number of predicted clusters per ground-truth pocket (fragmentation), while the **right diagram** shows the average number of ground-truth pockets per predicted cluster (merging). In an ideal one-to-one mapping, both means equal 1.0. While methods like DBSCAN reached high DCC recall, they showed excessive fragmentation (mean > 4). Our approach reduces this to 1.9.

| Binding site type | Method | Train set | $DCC_{top\text{-}N}$ | $DCC_{top\text{-}(N+2)}$ | $DCC_{MAX}$ |
|---|---|---|---|---|---|
| Small ($<$ 10 residues) | Seq2Pocket | sc-$PDB_{enhanced}$ | **50.8%** | **59.3%** | **63.6%** |
| | Seq2Pocket | sc-PDB | 41.7% | 48.3% | 50.8% |
| Large ($\geq$ 10 residues) | Seq2Pocket | sc-$PDB_{enhanced}$ | **74.6%** | **82.0%** | **84.6%** |
| | Seq2Pocket | sc-PDB | 67.3% | 74.1% | 76.0% |

**Table S1. Effect of data enhancement on small and large binding sites. To test the impact of training data enhancement, binding pockets in the LIGYSIS dataset were divided into two groups: *small* pockets containing fewer than 10 residues and *large* pockets containing 10 or more residues. DCC-based metrics were recomputed for each subgroup. Across all DCC variants, the improvements associated with training on sc-$PDB_{enhanced}$ are more pronounced for small binding sites.**

| Evaluation set | Method | Train set | $DCC_{top\text{-}N}$ | $DCC_{top\text{-}(N+2)}$ | $DCC_{MAX}$ | % RRO |
|---|---|---|---|---|---|---|
| CryptoBench (CBS) | Seq2Pocket 650M | CryptoBench train set | 64.7% | 73.0% | 74.0% | 49.6% |
| | *Seq2Pocket* | *CryptoBench train set* | *66.1%* | *76.3%* | *76.7%* | *52.9%* |
| LIGYSIS (GBS) | Seq2Pocket 650M | sc-$PDB_{enhanced}$ | 59.1% | 64.8% | 66.0% | 51.8% |
| | *Seq2Pocket* | *sc-$PDB_{enhanced}$* | *61.0%* | *68.8%* | *72.3%* | *52.9%* |

**Table S2. Finetuning the ESM-2 (650M) model. To improve accessibility for settings without computational resources required for large models such as ESM-2 (3B), we repeated the finetuning procedure using smaller ESM-2 (650M) variant. These models exhibit a modest decrease in pocket-level performance compared to their 3B counterparts.**

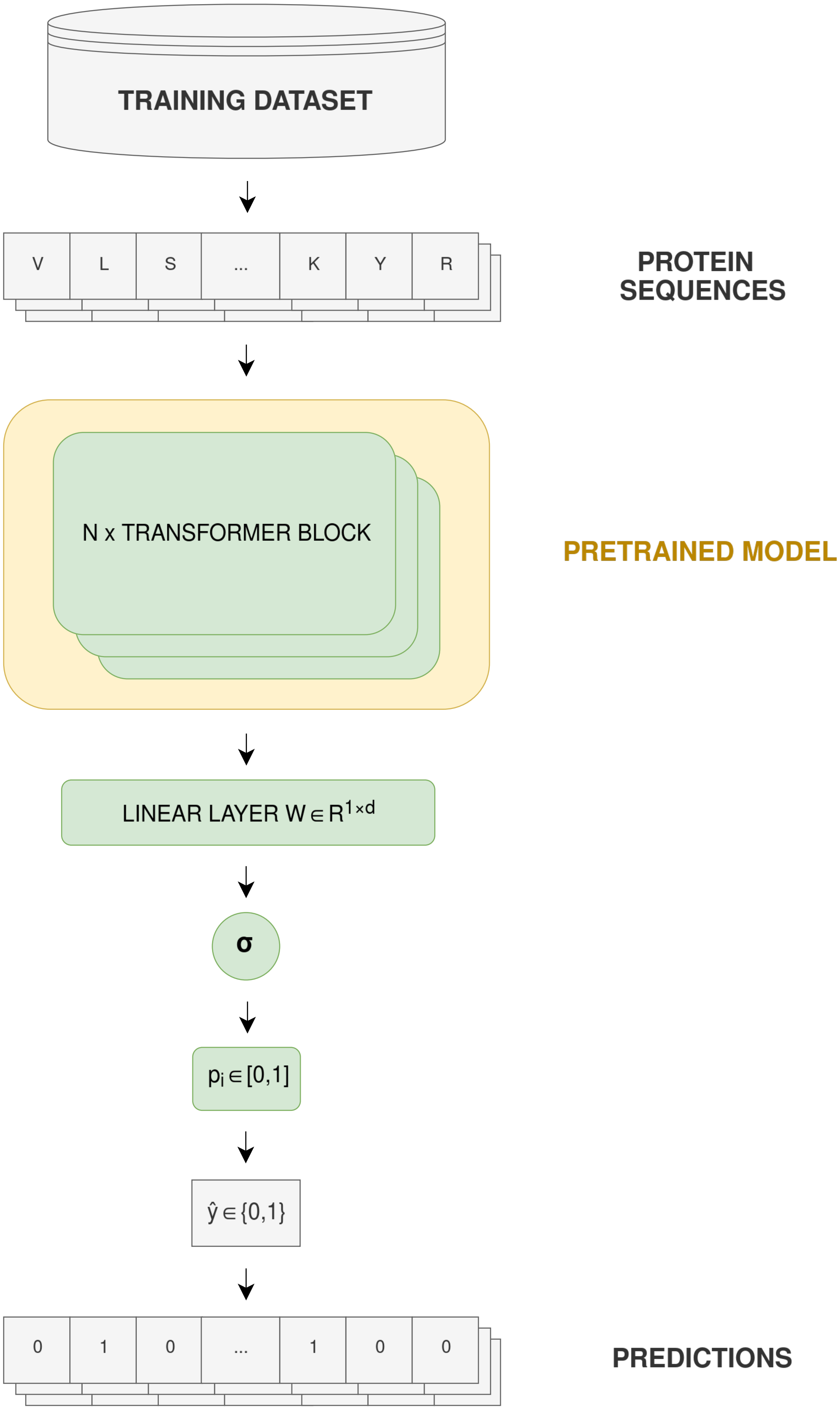


**Fig. S5. Protein language model (pLM) finetuning scheme.** During finetuning, the pLM (ESM2-3B) receives sequences from the training dataset - here either CryptoBench (2) for CBS prediction or sc-PDB (3, 4). for GBS prediction. Next, the original classification head is removed and replaced by a binary classification head for binding site prediction, see Linear Layer. Subsequently, the output of this layer is passed to a Sigmoid function ($\sigma$). This produces a probability $p_i \in [0, 1]$, which is binarized using a decision threshold into a single binary prediction for each residue in the sequence. Here, decision threshold was set to 0.7.

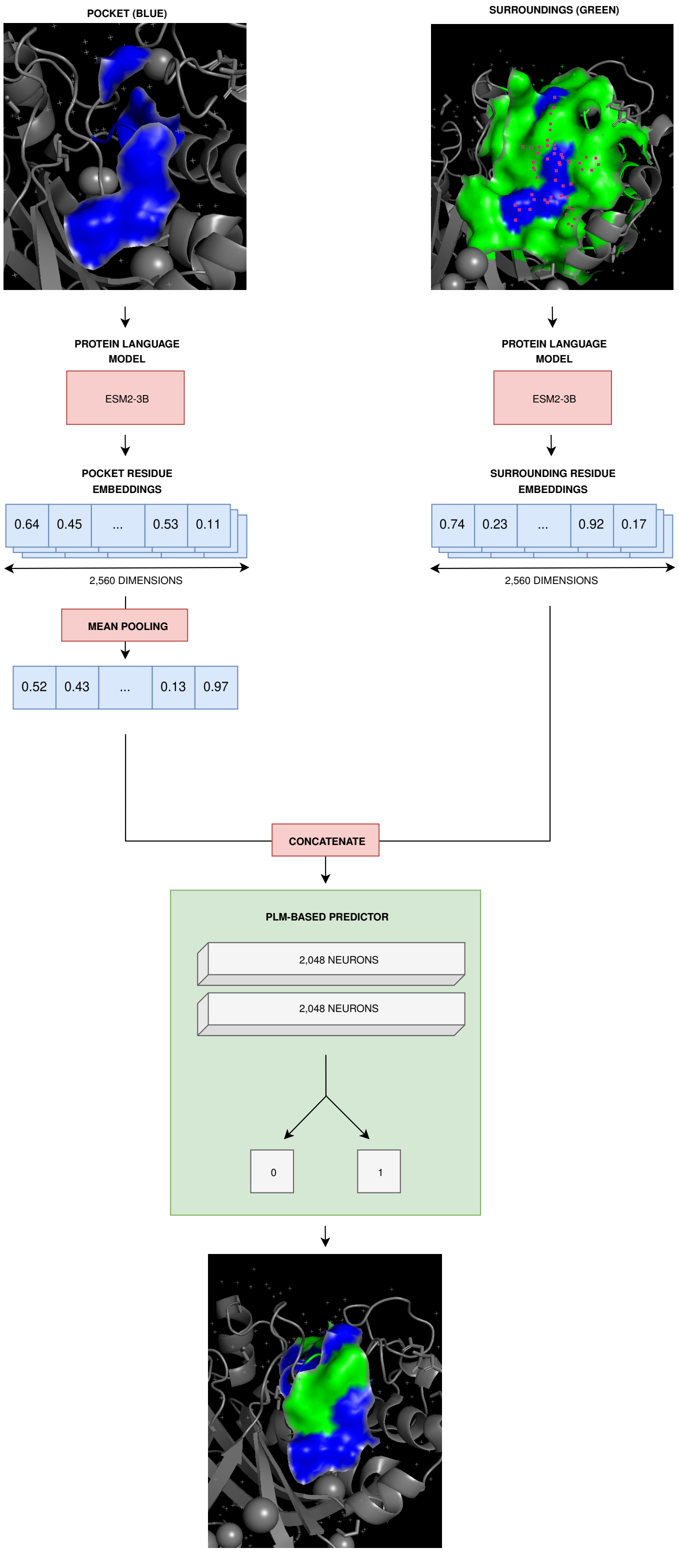


**Fig. S6. Architecture of the smoothing classifier.** The classifier processes two distinct inputs: (1) a global pocket representation, calculated as the mean average of all initial pocket residue embeddings (blue regions in the top-left and top-right picture), and (2) individual embeddings for all candidate residues located in the proximity of the predicted pocket (green region in the top-right picture). For each candidate, the model outputs a probability $p_i \in [0, 1]$ representing the likelihood of that residue being part of the pocket surface. These probabilities are binarized using a decision threshold of 0.4 to determine the final pocket membership. Final pocket is depicted on the bottom picture, where blue denotes the originally predicted residues, and green depicts the newly added residues.

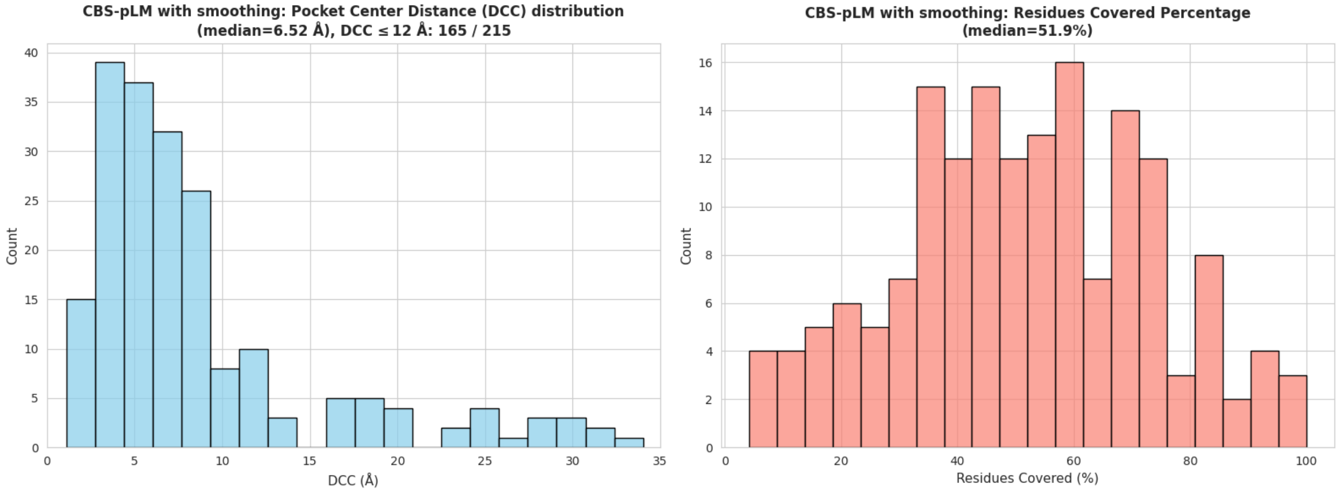


**(a)** Performance of CBS-finetuned pLM on the CryptoBench test set.

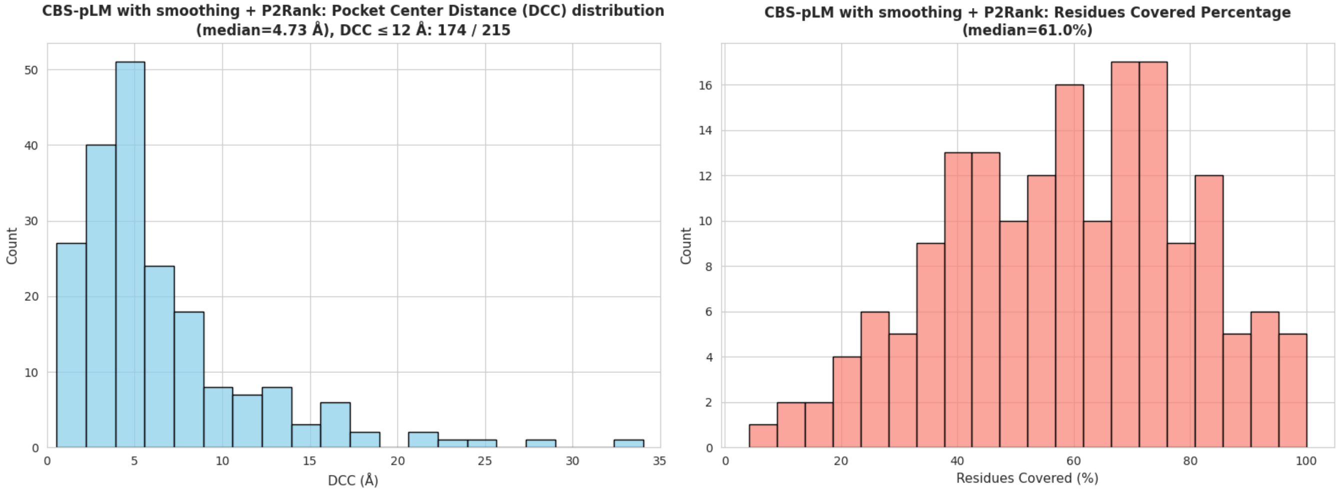


**(b)** Performance of integrated pLM and P2Rank (5) predictions on the CryptoBench test set.

**Fig. S7. Performance impact of integrating P2Rank with CBS-finetuned pLM predictions.** The left histogram displays the distribution of $DCC_{MAX}$, and the right histogram shows the distribution of RRO. Integration of P2Rank results with pLM predictions improves performance, resulting in a 4% increase in $DCC_{MAX}$ and a 9% improvement in *median* RRO.
